# Ten *Ostreobium* (Ulvophyceae) strains from Great Barrier Reef corals as a resource for algal endolith biology and genomics

**DOI:** 10.1101/2021.08.16.453452

**Authors:** Marisa M. Pasella, Ming-Fen Eileen Lee, Vanessa R. Marcelino, Anusuya Willis, Heroen Verbruggen

## Abstract

*Ostreobium* is a genus of siphonous green algae that lives as an endolith in carbonate substrates under extremely limited light conditions and has recently been gaining attention due to its roles in reef carbonate budgets and its association with reef corals. Knowledge about this genus remains fairly limited due to the scarcity of strains available for physiological studies. Here, we report on 10 strains of *Ostreobium* isolated from coral skeletons from the Great Barrier Reef. Phenotypic diversity showed differences in the gross morphology and in few structures. Phylogenetic analyses of the *tuf*A and *rbc*L put the strains in the context of the lineages identified previously through environmental sequencing. The chloroplast genomes of our strains are all around 80k bp in length and show that genome structure is highly conserved, with only a few insertions (some containing putative protein-coding genes) differing between the strains. The addition of these strains from the Great Barrier Reef to our toolkit will help develop *Ostreobium* as a model species for endolithic growth, low-light photosynthesis and coral-algal associations.

## INTRODUCTION

Corals are the results of a symbiotic association between animals, algae, and prokaryotes. The zooxanthellae – dinoflagellates in the family Symbiodiniaceae – colonize the coral tissue and provide the coral with its main source of carbohydrates. Over the years, many strains of the Symbiodiniaceae have been isolated, leading to major advances in our understanding of their taxonomy, photosynthesis, cell composition and interactions with the coral host (e.g. Hill *et al*. 2012, Goyen *et al*. 2015, LaJeunesse *et al*. 2018, Tortorelli *et al*. 2020).

In the calcium carbonate skeleton beneath the coral tissue, a green layer containing the green alga *Ostreobium* can often be seen. *Ostreobium* is the second major photosynthetic organism in the coral holobiont, with its biomass often exceeding that of Symbiodiniaceae (Odum & Odum 1955). Due to its peculiar niche burrowing into limestone substrates, studies of *Ostreobium* sp. have been few in comparison with Symbiodiniaceae.

Most of the physiological work done on *Ostreobium* used *in situ* measurements of pieces of coral skeleton rather than mono-algal culture strains. They highlighted how *Ostreobium* is shade-adapted and that some of the carbohydrates it produces through photosynthesis are transferred to the coral (Fine & Loya 2002). A handful of studies have used cultured strains, showing that *Ostreobium* is able to utilize a light absorption spectrum beyond 700 nm (near-infra-red, Fork & Larkum 1989, Koehne *et al*. 1999, Wilhelm & Jakob 2006).

During the last few years, there has been renewed interest in the genus, including work on its organelle and nuclear genomes (Verbruggen *et al*. 2017, Repetti *et al*. 2020, Iha *et al*. 2021). Although knowledge about the genus is steadily increasing, most of the studies have used a single *Ostreobium* strain, limiting our ability to generalize conclusions across the entire species complex. This is relevant because even though only a handful of *Ostreobium* species are formally described, environmental sequencing has shown that the *Ostreobium* clade is old and diverse, originating 500 million years ago and containing at least 80 species-level operational taxonomic units (OTUs) (Marcelino & Verbruggen 2016, Sauvage *et al*. 2016). Recent work shows that at least some of these *Ostreobium* OTUs differ in their physiology (Massé *et al*. 2020, Iha *et al*. 2021), illustrating the importance of having representative strains of different lineages to understand the breadth of physiological responses across the genus.

Until very recently, only two *Ostreobium* strains were available from public repositories (SAG strains 6.99 and 7.99). The former was isolated 30 years ago as an epiphyte on a red alga from the Philippines and the latter 20 years ago as an epiphyte of *Jania* sp. from southern Australia. In a recent paper, 9 closely related strains isolated from the coral *Pocillopora acuta* from the Aquarium Tropical du Palais de la Porte Dorée (Paris, France; originally from Indonesia) were deposited in the RBCell collection (Biological Resources of Living and Cryopreserved Cells) at the Muséum National d’Histoire Naturelle (MNHN; Paris, France; Massé *et al*. 2020).

The scarcity of available *Ostreobium* strains slows progress in understanding the biology of this genus and its interactions with coral and functions in reef decalcification. Here, we present a collection of 10 *Ostreobium* strains obtained from the skeletons of corals from the Great Barrier Reef, deposited in the Australian National Algal Culture Collection (ANACC, Hobart). Our aims for the paper are to describe the collection, isolation, and culturing procedures of these strains, provide their phylogenetic context and describe and compare their completely sequenced chloroplast genomes.

## METHODS

Coral fragments were collected with hammer and chisel at different sites and depths at Heron Island (Great Barrier Reef; Fig.1 and Table 1) from 8 colonies of *Porites* sp., 1 colony of *Pavona* sp., and one from an unidentified coral. From the green area in the skeleton, fragments of ca. 0.1 cm^3^ of volume were isolated with pliers and inoculated in 75mL culturing flasks at 26°C in f/2 medium (Guillard & Ryther 1962) but with vitamins provided at f/4 concentration (Guillard 1975). pH was kept at 8.1 ± 0.15 and salinity at 35ppm. Culturing flasks were transferred in a walk-in incubator under very low illumination: 1-2 μmol m^-2^s^-1^ of cool white LED light. When *Ostreobium* filaments started to emerge from the coral skeleton (Fig. 2), they were cut with a sterilized razor blade under an inverted light microscope to collect a single filament at a time. The newly collected filaments were transferred to a 24-well plate with the same medium used for the coral skeleton fragments. Once the filaments covered more than 50% of an individual well, they were transferred to plates with larger wells until enough biomass was reached for them to be moved to a 200mL glass culturing flask. Initially, some cultures were seen to contain diatoms, coccolithophores, prasinophytes, and cyanobacteria that were initially kept at very low levels by subculturing. At that stage, we avoided using antibiotics to allow *Ostreobium* to retain more of its natural microbiome for an ongoing research project.

**Figure 1-5.**
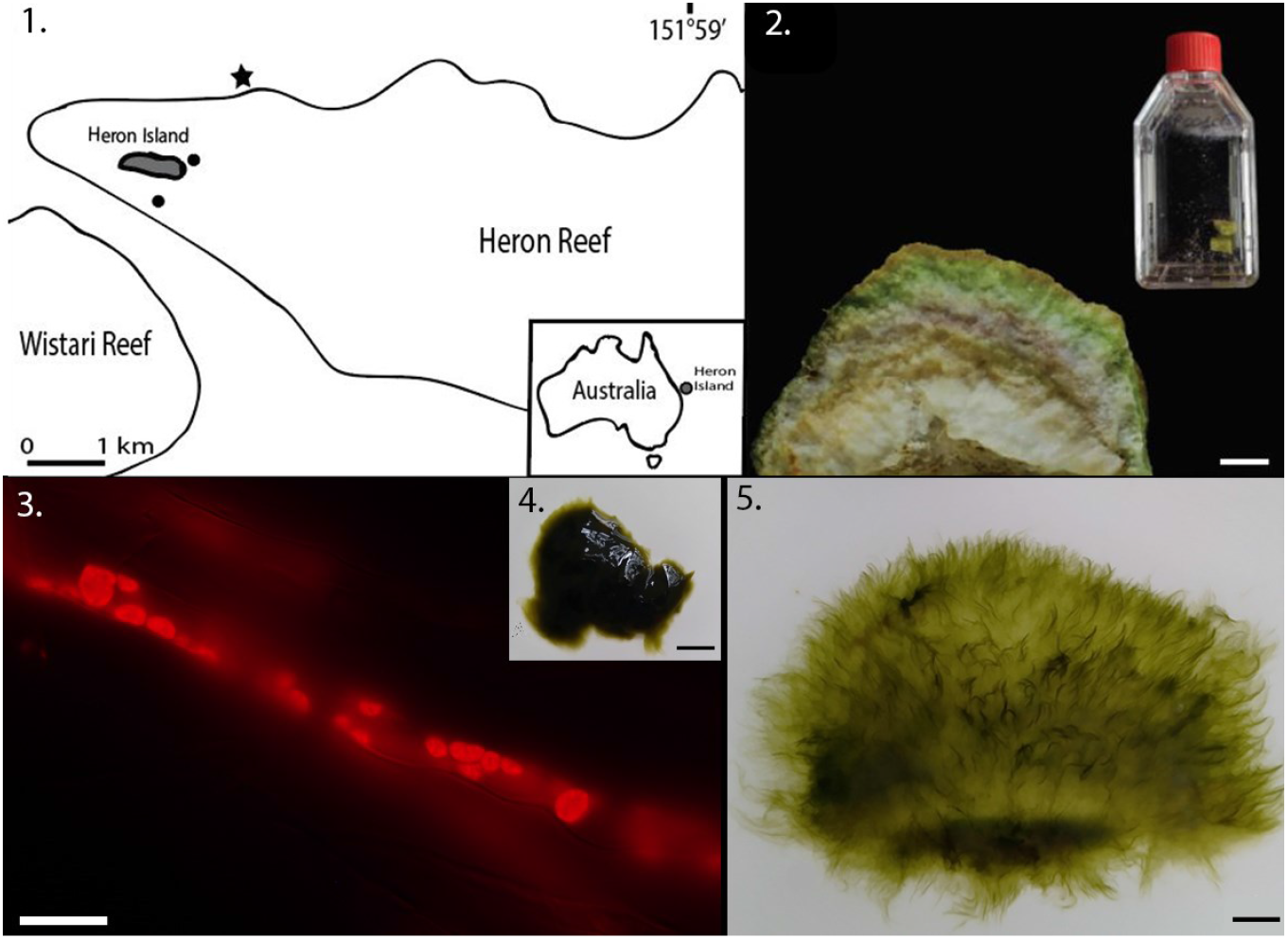
**1.**Sampling locations of the corals in Heron Island **2.** Longitudinal section of a massive coral skeleton showing the green layer of *Ostreobium* (scale bar 1 cm) and culturing vial containing coral fragments from which *Ostreobium* filaments are isolated. **3.** Filament of *Ostreobium* sp. imaged with fluorescence microscopy (scale bar 20μm). **4.** Compact thallus of free-living strain VRM650 (scale bar 1 cm). **5.** Diffuse morphology of VRM623 (scale bar 1 cm).

**Table 1.** Metadata for *Ostreobium* sp. strains isolated in this study. Strain VRM647 died during the COVID-19 lockdown and is not available in ANACC but is included here to provide the collection details and molecular data generated for this study.

| Strain | ANACC accession | Geographic Origin | Depth (m) | Longitude | Latitude | Phylogenetic affiliation |  | Coral Taxa | GenBank Accession |
| --- | --- | --- | --- | --- | --- | --- | --- | --- | --- |
|  |  |  |  |  |  | *rbcL* | *tufA* |  |  |
| VRM605 | CS-1379 | Heron Island (Research station beach) | 0.2-3 | 151.911965 | -23.4435 | CLADE P3/P14 | Lineage 3 | *Porites* sp. | XXXXXX |
| VRM609 | CS-1380 | Heron Island (Research station beach) | 0.2-3 | 151.911965 | -23.4435 | CLADE P3/P14 | Lineage 3 | *Porites* sp. | XXXXXX |
| VRM623 | CS-1381 | Heron Island (Shark bay) | 0.2-3 | 151.919524 | -23.4422 | CLADE P3/P14 | Lineage 3 | *Porites* sp. | XXXXXX |
| VRM627 | CS-1382 | Heron Island (Shark bay) | 0.2-3 | 151.919524 | -23.4422 | CLADE P3/P14 | Lineage 3 | *Porites* sp. | XXXXXX |
| VRM633 | CS-1383 | Heron Island (Shark bay) | 0.2-3 | 151.919524 | -23.4422 | CLADE P3/P14 | Lineage 3 | *Porites* sp. | XXXXXX |
| VRM638 | CS-1384 | Heron Island (Tenements) | 6 | 151.929604 | -23.4339 | - | - | - | XXXXXX |
| VRM642 | CS-1385 | Heron Island (Tenements) | 18 | 151.929604 | -23.4339 | CLADE C | Lineage 4 | *Porites* sp. | XXXXXX |
| VRM644 | CS-1386 | Heron Island (Tenements) | 16 | 151.929604 | -23.4339 | CLADE P1/K | Lineage 3 | *Unidentified coral* | XXXXXX |
| VRM646 | CS-1387 | Heron Island (Tenements) | 18 | 151.929604 | -23.4339 | CLADE P4 | Lineage 4 | *Pavona* sp. | XXXXXX |
| VRM647 | - | Heron Island (Tenements) | 10 | 151.929604 | -23.4339 | CLADE B3 | Lineage 4 | *Porites* sp. | XXXXXX |
| VRM650 | CS-1388 | Heron Island (Tenements) | 16 | 151.929604 | -23.4339 | CLADE B3 | Lineage 4 | *Porites* sp. | XXXXXX |

Substrains were also processed to obtain unialgal axenic strains at the Australian National Algae Culture Collection (ANACC). The ten *Ostreobium* substrains were maintained in 50mL f/2 medium in 200mL Petri dishes, at 20°C, under 1 μmol m^-2^s^-1^ photons LED light on a 12h:12h light: dark cycle. A small section of each strain was transferred to a 2mL Eppendorf tube containing 1mL f/2 containing an antibiotic cocktail (Penicillin 100 mg L^-1^, Streptomycin 25 mg L^-1^, Neomycin 25 mg L^-1^, Kanamycin 75 mg L^-1^) and incubated on a rotary mixer (Ratek, Australia) for 72h. The biomass was collected with tweezers, washed in fresh f/2 and placed in 4 mL f/2 in 6-well plates. Once biomass had increased, subsamples were transferred to f/2 agar plates, and subsequently, emerging filaments were cut off and transferred to 12-well plates in 2 mL f/2. Light microscopy and fluorescence microscopy (Vert.A1 Axio, Zeiss, Germany) with NucBlue DNA stain (Molecular Probes, Life Technologies, USA) was used to determine contamination of each strain, and the above procedure was repeated as needed to ensure cultures with no visible bacteria.

Pictures of each strain were taken with a CANON EOS 600D to characterize the gross morphology. Subsequently, autofluorescence images were taken using LAS X Widefield Systems with DM6000 B upright microscope at 63x magnification at the University of Melbourne-Biosciences Microscope Facility. All pictures of the strains have been deposited in FigShare (https://doi.org/10.6084/m9.figshare.15022026.v1).

For molecular identification and chloroplast genome sequencing, total genomic DNA was extracted following Cremen *et al*. 2016 and sequenced on the NovaSeq platform (paired-end, 150 bp reads, ca. 20 Gb per specimen) at GeneWiz (Suzhou, China) from libraries prepared with the Illumina VAHTS Universal DNA kit.

Sequence assemblies were generated *de novo* with MEGAHIT 1.2.9 (Li *et al*. 2015), SPAdes 3.14.1 (Bankevich *et al*. 2012) and the seed-and-overlap-based techniques NOVOplasty 4.2 (Dierckxsens *et al*. 2017) and GetOrganelle 1.7.1a (Jin *et al*. 2018). Completeness and circularity of the chloroplast genomes were tested using NOVOplasty 4.2 (Dierckxsens *et al*. 2017) and manual gap closing was performed by mapping the raw reads against the contigs obtained from the different assemblers in Geneious Prime 2020.1.3 (https://www.geneious.com). GeSeq (Tillich *et al*. 2017), ARAGORN (Laslett & Canback 2004) and MFannot (Beck and Lang 2010) were used as annotation tools and their information combined and curated in Geneious. Open reading frames (ORFs) were identified following the protocol of (Pasella *et al*. 2019) and the annotated plastid genome sequences were submitted to GenBank (Table 1).

The phylogenetic affiliation of the strains was inferred using two different chloroplast-encoded molecular markers derived from the genomes: elongation factor Tu (*tuf*A) and RuBisCo large subunit (*rbc*L). To provide context, we added *tuf*A sequences of 79 operational taxonomic units (OTUs) identified as *Ostreobium* sp. (Marcelino & Verbruggen 2016) and 56 *tuf*A of other Bryopsidales lineages as outgroups (Table S1). For *rbc*L, we included 81 *rbc*L identified as *Ostreobium* sp., including the *rbc*L sequences of Gutner-Hoch & Fine 2011 and Massé *et al*. 2020 and 20 *rbcL* of other Bryopsidales (table S1). For both molecular markers, sequences were aligned using MUSCLE (Edgar 2004) in Geneious Prime 2020.1.3 (https://www.geneious.com) and the phylogenetic tree for *tuf*A was inferred with maximum likelihood using RAxML v8.0.26 (Stamatakis 2014). We used GTR + Γ as the model of sequence evolution and branch support was estimated using 100 bootstrap replicates.

## RESULTS & DISCUSSION

We initially isolated 11 strains of *Ostreobium* from coral skeleton fragments collected on Heron Island (Australia, GBR). Strain VRM647 died soon after we sequenced it, so ten unialgal strains were deposited in the Australian National Algal Culture Collection (ANACC, CSIRO, Hobart; Table 1). After several months of growth at low illumination, the free-living thalli showed apparent variation, with gross morphology varying from very compact, dark green thalli to relatively diffuse structures with filaments that appear less pigmented despite growing in identical conditions. Strain VRM650 (Fig. 4) presented very dark compact thalli, while strains VRM605, VRM609, VRM623, VRM633, VRM644 and VRM646 shared the same diffuse morphology (Fig. 5) while only presented and the remaining strains showed an intermediate filament density.

All strains are composed of undifferentiated cylindrical filaments with a diameter between 8 and 13 μm. No swellings or cross-walls were observed in any filaments. For strain VRM623, we found constrictions at random intervals as reported previously for *Ostreobium constrictum* (Lukas 1974). Kobara & Chihara (1992) reported “sporangia-type organs” in *Ostreobium*, occurring at the ends of some filaments. We also observed these in all of our strains, and we noticed new filament growth on the glass culture flasks for several strains. While we could not make microscopic observations to confirm whether these filaments grew from spores that were released from the sporangia-like structures, this is the most likely explanation for our observations.

Chloroplasts were not observed to be reticulated as reported by Lukas (1974). Instead, they were large and ovoid and often nearly as wide as the filaments for all the strains (Fig. 3). The chloroplasts appeared to be homogeneously distributed along all the filaments, in contrast to previous reports of free-living cultures of *Ostreobium*, where the chloroplasts were reported to be small and mainly distributed against the cytoplasmic side of the cell membrane (Massé *et al*. 2020).

*Ostreobium* diversity is mostly known from environmental sequencing. Different studies have used either *tuf*A or *rbc*L as a marker gene, and as a consequence, two alternative classifications have been proposed (Gutner-Hoch & Fine 2011; Marcelino *et al*. 2016; Sauvage *et al*. 2016; Massé *et al*. 2020). Here, we provide phylogenies for the newly established strains of *Ostreobium* sp using both markers in order to place the strains in both systems.

The phylogeny based on the *tuf*A gene (Fig. 6) showed that the newly established strains belong to *Ostreobium* lineages 3 and 4, following the classification by Marcelino *et al*. 2016. All strains collected from the shallow waters of Shark Bay and Research Station Beach belong to lineage 3 in the *tuf*A phylogeny, while the strains from the deeper-water Tenements site are part of lineage 4, except for VRM644, which is in lineage 3. Among the strains in lineage 3, four strains (VRM605, VRM609, VRM627, and VRM633) have identical *tuf*A sequences, indicating that they are conspecific, and VRM623 is very closely related to them (98.5% similarity). In lineage 4, VRM650 and VRM647 are likely conspecific (99.2% similarity). The remaining strains are more distantly related and recovered in different positions in the phylogenetic tree (Fig. 6).

**Figure 6.**
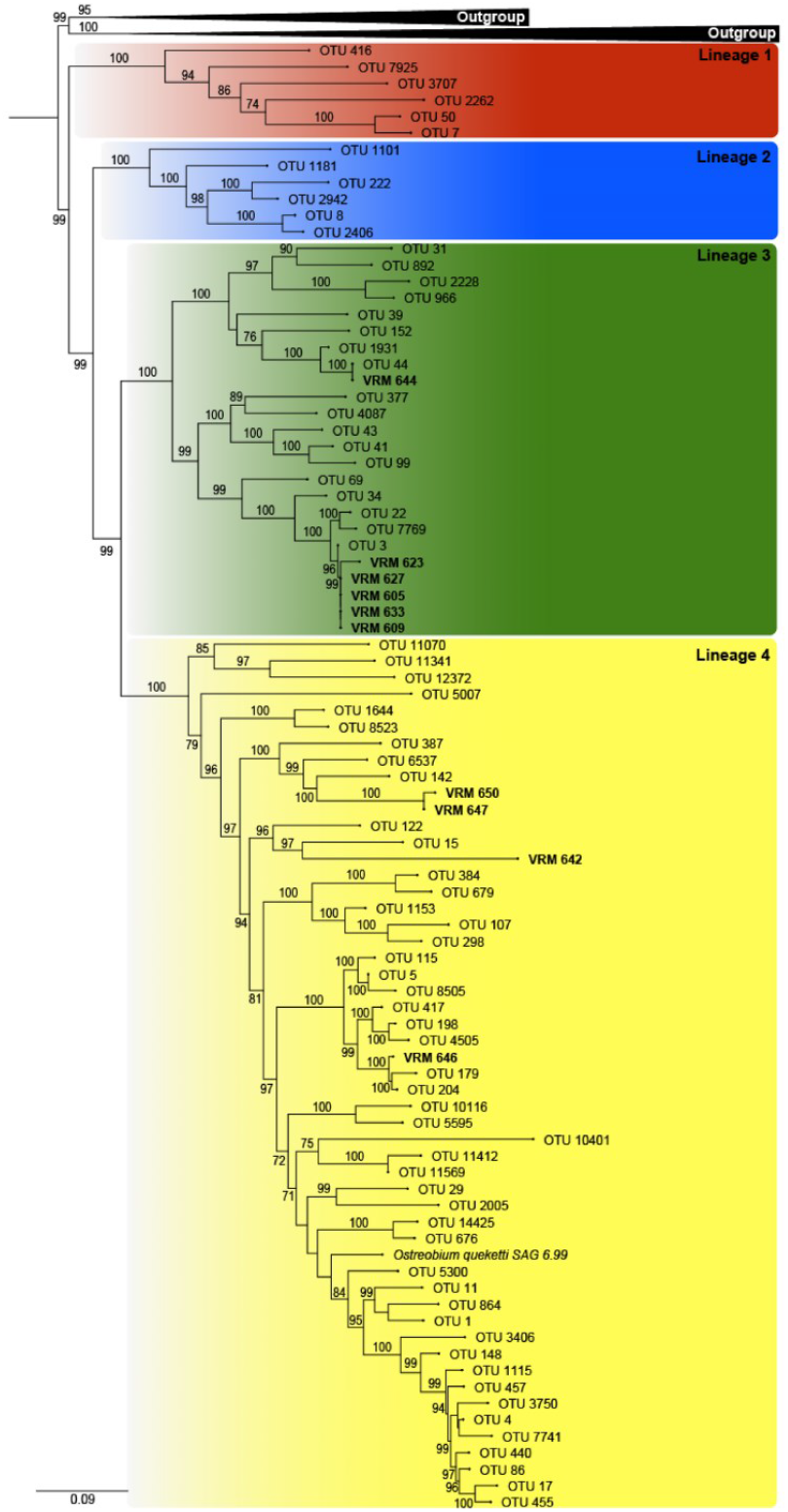
Maximum likelihood tree of *Ostreobium* lineages based on *tuf*A sequences. New strains isolated in this work are indicated in bold. Lineage numbering follows Marcelino *et al*. (2016). Bootstrap values above 70 are shown. The GenBank accession numbers are listed in Table 1 and Table S1. Note that the phylogeny also includes VRM647, an isolate for which we have molecular data available but that has died.

The strains in the *rbc*L tree were annotated with the classification provided by Massé *et al*. 2018 (Fig. 1S). As was the case for *tuf*A, the *rbc*L phylogeny showed strains from Shark Bay and the Research Beach Station belonging to a single clade, P3. The remaining five strains belong to different clades (Table 1; Fig. 1S). Strain VRM644 is most closely related to the P1-clade strains isolated by Massé *et al*. (2020).

We obtained the complete chloroplast genomes and confirmed their circularity for all the strains except VRM623 (Table S2). The size and the GC content of the newly sequenced *Ostreobium* chloroplast genomes are similar to those reported previously (Marcelino *et al*. 2016, Del Campo *et al*. 2017, Verbruggen *et al*. 2017). A total of three ribosomal RNAs and 77 chloroplast protein-coding genes were shared by all the newly assembled chloroplast genomes. All strains encoded 31 tRNA genes, of which tRNA-Met was present in 3 copies as previously reported for this alga (Marcelino *et al*. 2016; Verbruggen *et al*. 2017). The large ribosomal protein L19 (*rpl*19) gene was lost in four closely-related *Ostreobium* strains in the lineage 3 (Table S2). The ribosomal protein L19 contributes to bridging the two ribosomal subunits in bacteria (Gao *et al*. 2003) and it is not currently clear whether the loss observed in these *Ostreobium* strains represents a transfer to the nucleus or a genuine loss. The gene has been lost from the plastid genome on several occasions in the evolution of green-type algae (Uthanumallian *et al*. 2021), but this is the first time that it was shown to be lost from the plastid in the order Bryopsidales.

All the chloroplast genomes appeared to have only slight variations in the genome size, except for the strain VRM644 that presented two fairly large insertions, one being a 1.8 kb insertion between the *rpl*32 and *rps*9 genes and the second a 1.3 kb insertion between the *psb*H and *clp*P genes. In each insertion, we found a freestanding (i.e. non-intronic) ORF (*orf*237 and *orf*100) but BLAST searches did not identify any similarity with previously annotated protein-coding genes in green algal chloroplast genomes. Similarly, searches of the amino acid sequences in the NCBI conserved domain search did not return any results. So, even though the origins and functions of many ORFs in the plastid genomes of Bryopsidales remains unclear, recent transcriptomic work showed that many of them are expressed (Zou *et al*. 2021), suggesting that they do serve a function in the cell.

Three others freestanding ORFs were found in other *Ostreobium* chloroplast genomes, of which two were exclusive to VRM642, inserted between *chl*B and *psa*A (*orf*122 and *orf*134; table S2), and one was found in all the strains between the genes *psb*A and *chl*I. It showed significant sequence similarity to a previously reported freestanding ORF encoding a group II intron RT/maturase in Bryopsidales. The freestanding group II intron RT/maturase in the Ostreobineae is hypothesized to play a key role in promoting slicing of the introns, since this lineage does not encode genes that may facilitate splicing of its group II introns (Cremen *et al*. 2018).

In all the newly sequenced chloroplast genomes, we found group II introns in *rpo*B, *rpl*23, *rpl*5 and *rpo*C1, as had previously been reported for other *Ostreobium* strains (Marcelino *et al*. 2016; Verbruggen *et al*. 2017). However, the presence of introns in other genes was strain-specific (Table S3). Group II introns were also found in *rps*4 and *ycf*3 for strains VRM650, VRM647, VRM646 and VRM642.

The tRNA ile-lysidine synthetase (*til*S) presents four different types of gene fragmentation in the Bryopsidales (Cremen *et al*. 2018). In the newly sequenced *Ostreobium* chloroplast genomes, seven strains showed fragmentation of the gene with a frameshift, and VRM646 having both an in-frame stop codon and a frameshift. The gene *til*S of the strain VRM642 was the only one not presenting any type of fragmentation, as was already reported for a previously sequenced *Ostreobium* chloroplast genome (Del Campo *et al*. 2016).

## CONCLUSION

The isolation of 10 *Ostreobium* strains from skeleton fragments of Great Barrier Reef corals offers perspectives for expanding our knowledge of the biology of this genus. Along with previous strains, they are a valuable addition to our toolkit for this emerging algal model for low-light photosynthesis, endolithic biology and coral symbiosis research. Our work provided detailed information about their phylogenetic context using both commonly used genetic markers and illustrates variations in their chloroplast genomes.

## ACKNOWLEDGEMENTS AND FUNDINGS

We thank Trevor Bringloe for logistical assistance and Allison Van de Meene of the University of Melbourne - Biosciences Microscopy Facility for help with microscopy. Authors also thank the Heron Island Research Station staff for the support provided during the sampling campaign. Funding was provided by the Australian Research Council (DP200101613 to HV) and samples were collected under permit G13/36490.1.

## SUPPLEMENTARY MATERIALS

**Figure S1.**
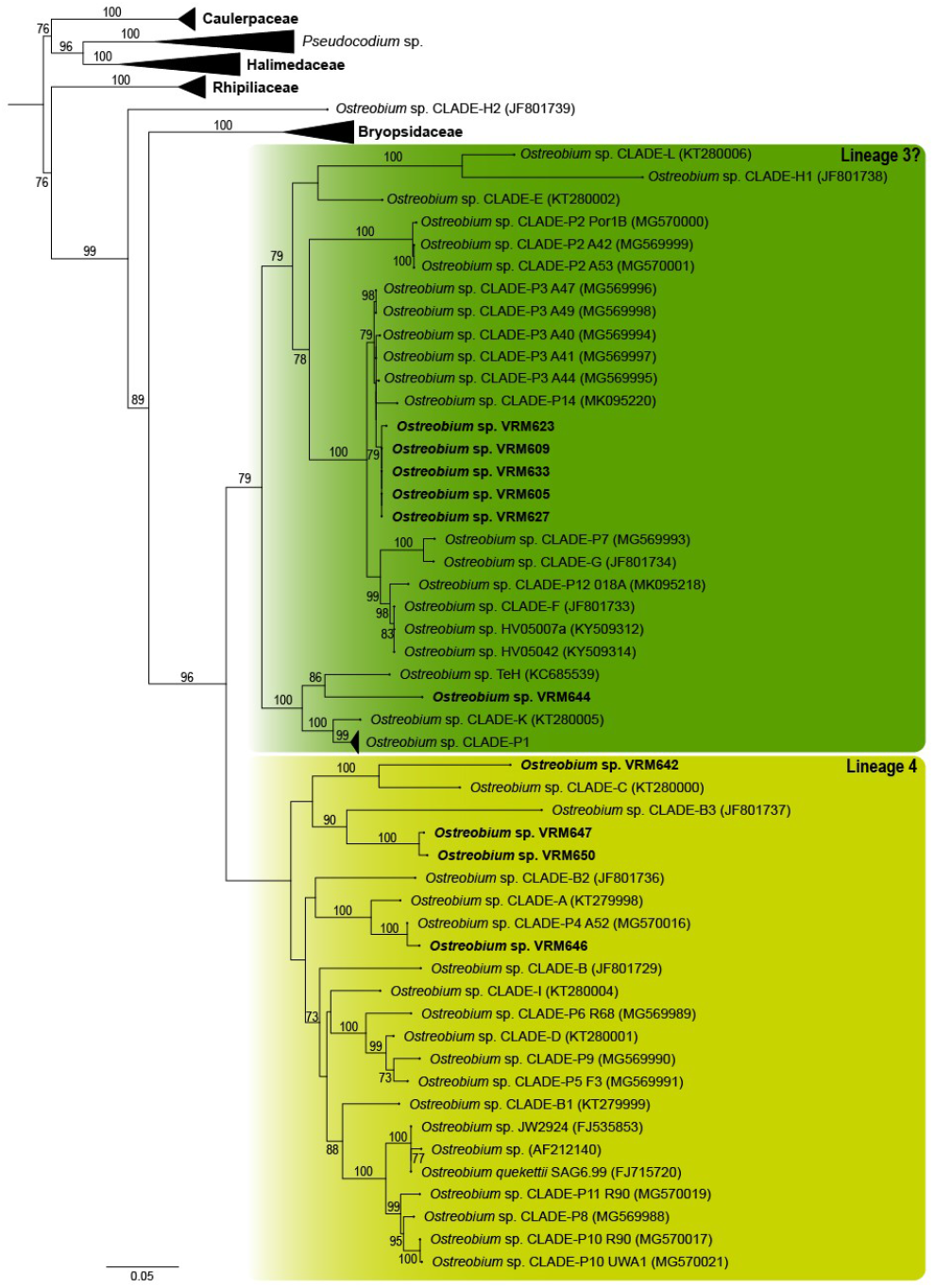
Maximum likelihood tree of *rcb*L sequences. Specimens in boldface are the strains isolated in this work. The taxon labels indicate the *rbc*L clade names from Massé *et al*. (2018). The colours show how we think the major lineages 3&4 from the *tuf*A tree map onto the *rbc*L tree. GenBank accessions of new sequences are listed in Table 1 and Table S1. Only bootstrap values above 70 are shown.

**Table S1.**

Reference sequences Genbank accession number used to reconstruct the green algae tufA phylogeny and outgroup of the rbcL phylogeny.

**Table S2.**

Features of the chloroplast genome in the *Ostreobium* strains.

**Table S3.**

Comparison of the Group II introns distribution in the genes of the *Ostreobium* sp strains.

